# Canopy height model and NAIP imagery pairs across CONUS

**DOI:** 10.1101/2024.12.24.630202

**Authors:** Brady W Allred, Sarah E. McCord, Scott L. Morford

## Abstract

Canopy height models (CHM) provide detailed environmental vertical structure information and are an important indicator and input for ecological and geospatial applications. These models are often spatiotemporally inconsistent, necessitating additional modeling to scale them in space and time. Yet, such scaling is hindered by a lack of spatially diverse data. To address this, we use United States Geological Survey 3D Elevation Program lidar data to produce 22,796,764 one meter resolution CHM chips, stratified across the dominant land covers of the conterminous United States. For each CHM, we pair a matching time-aligned aerial image from the United States Department of Agriculture National Agriculture Imagery Program. This dataset can be used to train models for large scale CHM production.

## Background and Summary

Canopy height models (CHM) are spatially explicit representations of the vertical structure of an environment, measured relative to the ground surface. These models provide detailed information about the structure, arrangement, and organization of vegetation and the built up environment; they are used in numerous applications, including land management and conservation, carbon and climate change modeling, landscape and habitat monitoring, disaster risk assessment and management, and geospatial analysis and modeling^1–5^. As CHMs are commonly derived from airborne lidar (i.e., Light Detection and Ranging) instruments, the availability of CHMs are often limited to local or regional acquisitions and to single snapshots in time. Spaceborne lidar (e.g., GEDI and ICESat-2) provide increased and often repeated coverage, but with the tradeoff of coarser ground sampling distances.

To overcome these challenges and produce CHMs at scale, recent approaches have combined lidar derived CHMs with multispectral optical or radar imagery. Combining spaceborne GEDI derived CHM with Landsat, Sentinel-2 or Sentinel-1 has resulted in modeled estimates of canopy height at global and regional scales^6–9^. Aerial lidar derived CHMs have also been successfully utilized, most commonly with high resolution aerial or satellite imagery. Wagner et al.^10^ produced sub-meter canopy height estimates for the state of California by producing their own aerial derived CHMs and training a model with United States Department of Agriculture (USDA) National Agriculture Imagery Program (NAIP) images. Notably, Tolan et al.^11^ produced sub-meter global estimates of canopy height through self supervised learning of Maxar satellite imagery and subsequent training of off-the-shelf CHMs (both air and spaceborne).

Due to the ability of tree height to reduce uncertainty in woody plant carbon modeling, scientists have focused CHM development and modeling efforts within forested ecosystems. Other ecosystems are often neglected or excluded entirely from model training, reducing their accuracy and utility in these ecosystems. Rangelands (inclusive of grasslands, savannas, and shrublands) have received little attention with regard to CHM modeling efforts, even though they are a dominant land cover^12^ and canopy height measurements are valuable to numerous modeling pursuits and on-the-ground management^3^. The inherent heterogeneity of rangelands (i.e., a mixture of grasses, shrubs, trees, or lack thereof) often requires fine resolution CHMs, reducing uncertainty and making them more suitable for application^13^.

The paucity of high resolution, aerial derived CHMs presents difficulties when training models for broader scale application. In the United States, the United States Geological Survey (USGS) works with partners to collect aerial lidar for the 3D Elevation Program (3DEP)^14,15^. Although data are collected in different regions, at different times, by different contractors, and ultimately processed into various derived products (primarily digital elevation models), lidar data are publicly available for independent CHM production. The overhead of retrieving and processing these data, however, can be challenging.

We used the USGS 3DEP lidar collection to produce a geographically large, but spatially disparate, CHM dataset. We focused our efforts on United States rangelands, but ensured that other dominant land covers are included. Our dataset comprises 22,796,764 CHM images, each spatially paired with a USDA NAIP image.

## Methods

### Location sampling

Utilizing the availability of USGS 3DEP lidar data and USDA NAIP imagery, we focused dataset development within the conterminous United States (CONUS). We stratified location sampling by Environmental Protection Agency level three ecoregions and National Land Cover Database (NLCD; 2019 release) dominant classes. Land cover subclasses were aggregated to their dominant class (i.e., deciduous, evergreen, and mixed forest were aggregated to forest), with the exception of pasture and cultivated crops. Our baseline sampling was 50,000 locations of each class within an ecoregion. To ensure greater representation of rangelands, we increased sampling of herbaceous and shrubland classes by 4x and the pasture class by 2x. We decreased sampling of the water class by 0.1x to limit its abundance.

To maintain a minimum distance of 240m between sampling locations across all classes, we aggregated the NLCD to 240m resolution by calculating the mode of all pixels within the aggregation unit. This sampling produced approximately 30.5 million locations, which were further reduced by availability of lidar data and NAIP imagery.

### Lidar data

USGS 3DEP lidar data are available as LAZ format tiles via USGS rockyweb (https://rockyweb.usgs.gov/) and Amazon Web Services (AWS) cloud storage (https://registry.opendata.aws/usgs-lidar/). Additionally, many copies of the data are also available in Entwine Point Tile (EPT) format via AWS cloud storage. EPT format is cloud-friendly and streamable, allowing users to easily retrieve and process lidar based on geographic location or other parameters. We used the EPT format to construct this dataset.

USGS 3DEP lidar data are published by work units. Using the USGS Work unit Extent Spatial Metadata (WESM) we selected work units that met the following criteria: 1) lidar data collected from 2014 through 2023; 2) had a Quality Level (QL) of QL2 or lower (see Table 1); and 3) had a LPC category of “Meets”, “Meets with variance”, or “Expected to meet”, referencing the data’s ability to meet 3DEP specifications. We selected work units available in EPT format and buffered their perimeters inward by 200m to reduce edge effects. We selected sampling locations that intersected work units, thereby reducing sampling locations to approximately 23.2 million.

**Table 1.** USGS 3D Elevation Program (3DEP) quality requirements. Only data meeting QL2 or lower were used in this study. NVA references non-vegetated vertical accuracy; VVA references vertical vegetated accuracy. From Stoker and Miller ^14^.

| Quality Level | RMSE<sub>z</sub> non-vegetated (m) | NVA 95% confidence level (m) | VVA 95th percentile (m) | Aggregate nominal pulse spacing (m) | Aggregate nominal pulse density (pls/m<sup>2</sup>) |
| --- | --- | --- | --- | --- | --- |
| QL0 | ≤ 0.05 | ≤ 0.098 | ≤ 0.15 | ≤ 0.35 | ≥ 8.0 |
| QL1 | ≤ 0.10 | ≤ 0.196 | ≤ 0.30 | ≤ 0.35 | ≥ 8.0 |
| QL2 | ≤ 0.10 | ≤ 0.392 | ≤ 0.30 | ≤ 0.71 | ≥ 2.0 |

We used the Point Data Abstraction Library (PDAL) to access lidar data. For each sampling location, we buffered outward with a square radius of 200m and retrieved all lidar point data within this perimeter. Lidar point data were reprojected to the appropriate UTM zone of the sampling location and temporarily stored as LAZ files. Due to various errors, we were unable to retrieve lidar data for 156 sample locations.

### Canopy height model production

We used the lidR R package to produce CHMs. We excluded sampling locations where ground classifications were absent (i.e., densely vegetated areas). We classified points as noise using the ivf() algorithm with a voxel resolution of five meters and a maximum of six points. Height was normalized using k-nearest neighbor and inverse-distance weighting methods. We excluded points classified as noise and points where the normalized height was greater than 300m. We did not remove aboveground, non-vegetated data prior to normalization; accordingly, the produced CHMs are normalized digital surface models. As the vast majority of our sampling locations were in vegetated areas, we retained the use of the CHM label to describe our dataset.

We produced CHMs at a resolution of one meter using the pitfree() algorithm (an implementation of the algorithm developed by Khosravipour et al.^16^), with threshold parameters of 0, 2, 5, 10, 15, maximum edge parameters of 10, 1, and a subcircle parameter of 0.35. If CHM production failed with the pitfree() algorithm, we attempted to generate a CHM using the p2r() algorithm with a subcircle parameter of 0.35. We abandoned CHM production after any subsequent failures. The acquisition date was retrieved from the GPS time of the lidar data. We cropped CHMs to dimensions of 256×256 pixels (centered on the sampling location), scaled height values by 100, and wrote files as GeoTiffs. We could not produce CHMs for approximately 370,000 sample locations due to absence of ground classifications, noise, or other errors. Additionally, around 10,000 sample locations were removed because their acquisition dates were before 2014. We produced more than 22.8 million CHMs across CONUS.

### NAIP imagery retrieval

States collect USDA NAIP imagery at various intervals, typically every 2-3 years. Ground sampling distance (GSD) of NAIP varies from 0.3m to 1.0m, with 0.6m being the most common for images acquired since 2014. Each NAIP image includes four spectral bands: Red (R), Green (G), Blue (B), and Near-Infrared (N). For each CHM produced, we retrieved available NAIP images two years before and after the lidar acquisition date. As this four year window may contain multiple NAIP collection events (corresponding to the specific year of collection), we identified the collection event closest to the lidar acquisition date and selected images from this event only. This approach ensured that all NAIP images for a given CHM originated from the same collection event and shared the same GSD.

We mosaicked and cropped NAIP images to match the spatial footprint of the corresponding CHM. The GSD was retained from the original NAIP images, resulting in variability among sampling locations and years. We included all RGBN bands and added a fifth “mask” band to indicate pixel validity (0 for invalid pixels and 1 for valid pixels). We retrieved NAIP imagery from Google Earth Engine^17^. Approximately 90,000 sample locations had no NAIP imagery available during the specified four year time window.

In total, we produced 22,796,764 spatially matching CHM and NAIP pairs across CONUS.

### Data Records

CHM and NAIP pairs are available as a collection of tar files at http://rangeland.ntsg.umt.edu/data/rap/chm-naip/. The tar files are separated by CHM and NAIP, and organized hierarchically by 1) UTM zone; and 2) the first three digits of the y coordinate (UTM northing) of the sampling location. Total size of the dataset is approximately 12 TB. A CSV file is provided with the relative path of each pair; UTM zone, x and y coordinate of the sampling location; date of CHM and NAIP acquisition; and sampling location dominant land cover and ecoregion classification.

CHM GeoTiffs are height values in meters, scaled by 100, and stored as a single band 16-bit integer, with a no data value of 65535. NAIP GeoTiffs are stored as five bands (RGBN plus a “mask” band) 8-bit integer. A mask value of 0 indicates an invalid pixel; a value of 1 indicates a valid pixel.

### Technical Validation

The geographic distribution of CHM and NAIP pairs reflected the sampling design and data availability (Fig. 1). As intended, rangelands were oversampled relative to other land cover classes (Fig. 2A). Distributions of the acquisition date for CHM and NAIP images were similar, with many being acquired in years 2018 to 2020 (Fig. 2B,C). The average absolute difference between CHM and NAIP acquisition dates was 200 days. We note a slightly skewed distribution in the differences of acquisition dates between CHM and NAIP images, with more NAIP images acquired before CHM acquisition rather than after (Fig. 2D). This is likely due to the fact that 1) as we approach the end of the sampling period, only past images are available; and 2) there can be a significant lag in the publication of NAIP imagery.

**Figure 1.**
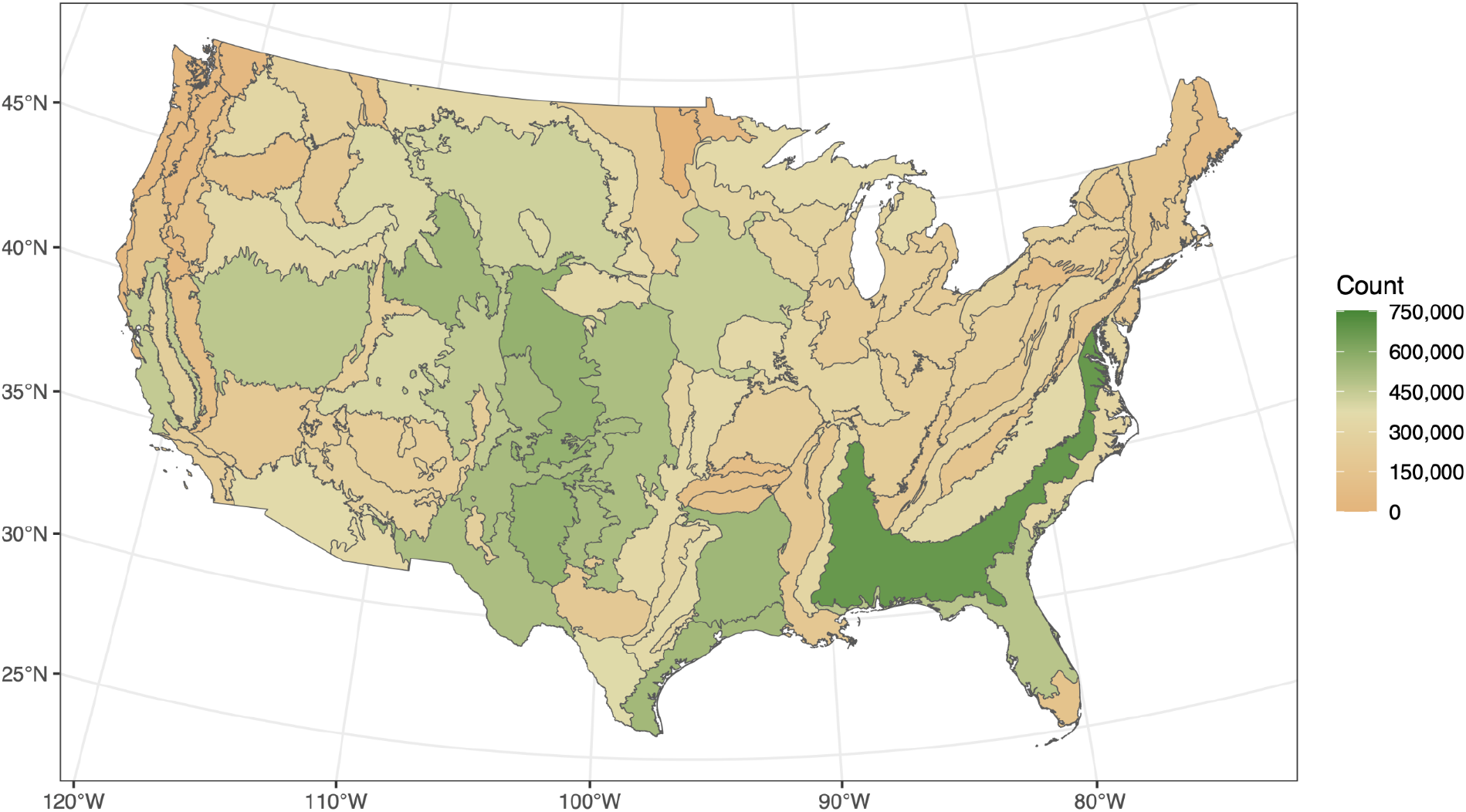
Geographic distribution of canopy height model and NAIP image pair locations by EPA Level III ecoregion. Canopy height model generation was limited by USGS 3DEP lidar data acquired from 2014 through 2023 and quality requirements.

**Figure 2.**
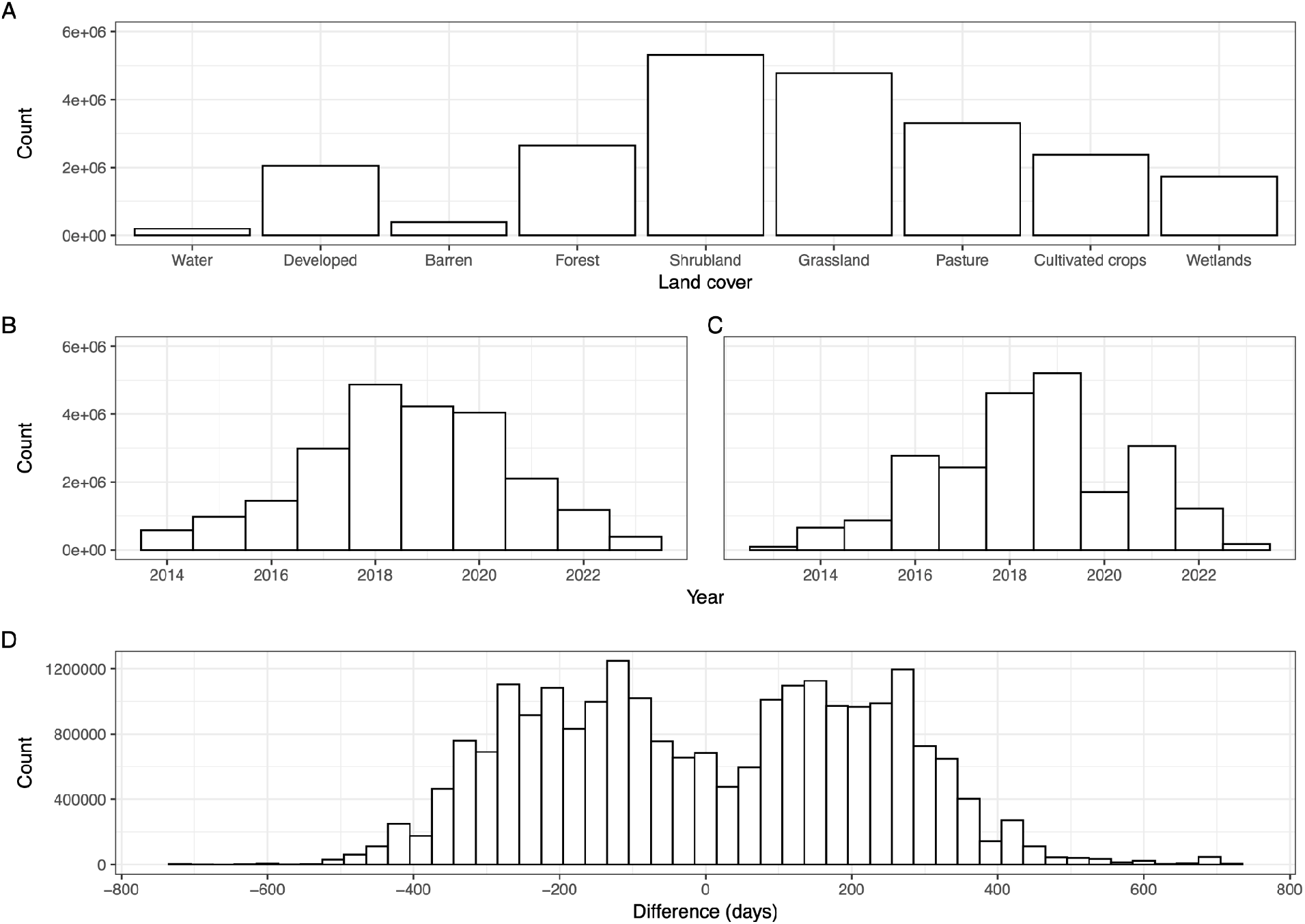
Dataset frequencies. A) Frequency of dominant land cover class of sampling locations. Shrubland, grassland, and pasture were intentionally oversampled; water was intentionally undersampled. B) Frequency of canopy height model year. C) Frequency of NAIP image year. D) Frequency of the time difference (days) between lidar acquisition and USDA NAIP for canopy height model and NAIP pairs. The closest NAIP collection event to the lidar acquisition date (within a four year window, two years before and two years after) was used to construct the matching NAIP image.

Our approach successfully paired NAIP imagery with CHMs (Figs. 3 and 4). Some NAIP images, however, were partially or completely empty due to their placement on the spatial edge of a collection event or to missing data (Fig. 5). Additionally, due to potential time differences between a CHM and NAIP image, significant landscape change may have occurred (Fig. 6).

**Figure 3.**
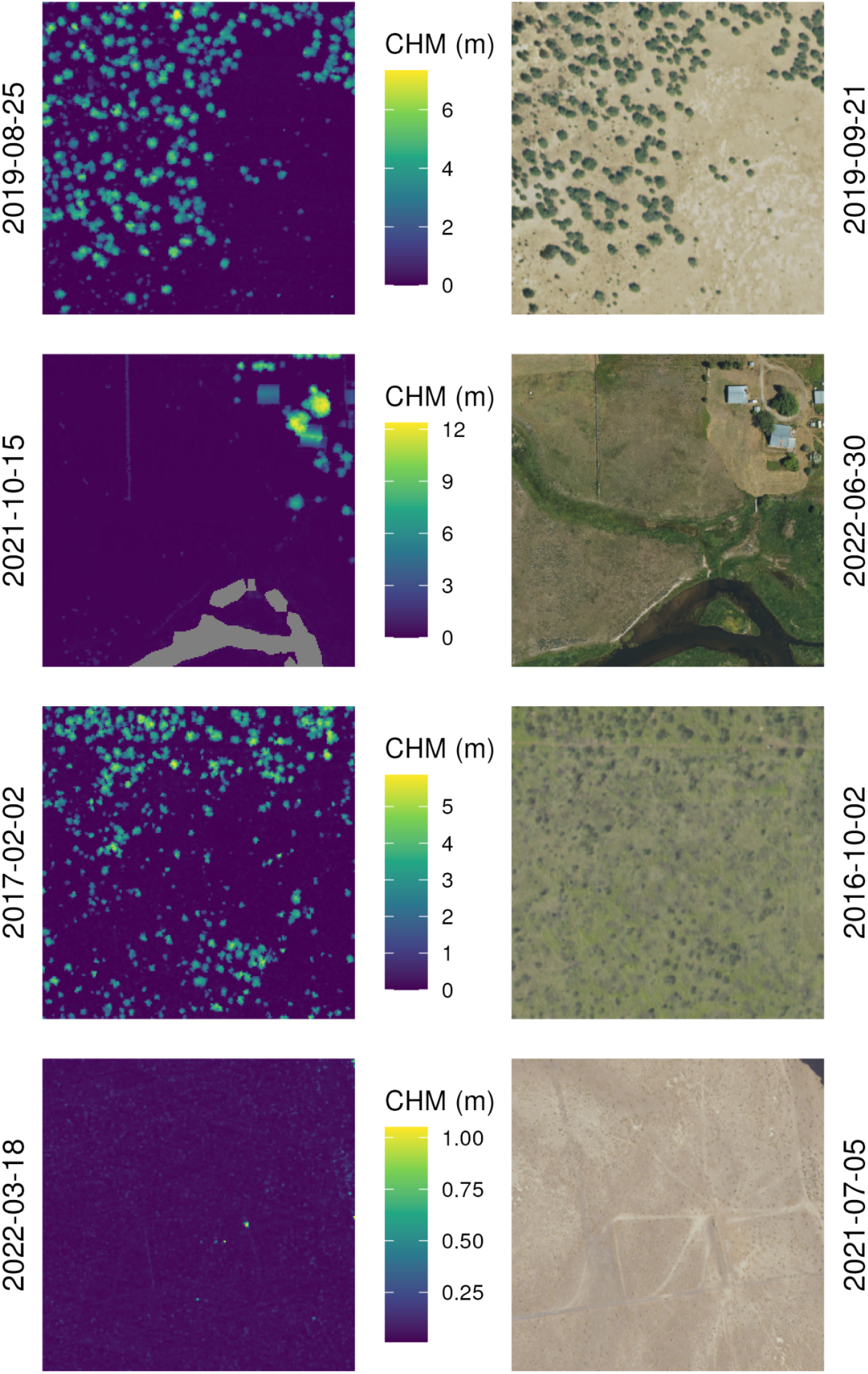
A sample of paired canopy height model (CHM; left) and NAIP images (right). Dates on vertical axes represent the date of acquisition.

**Figure 4.**
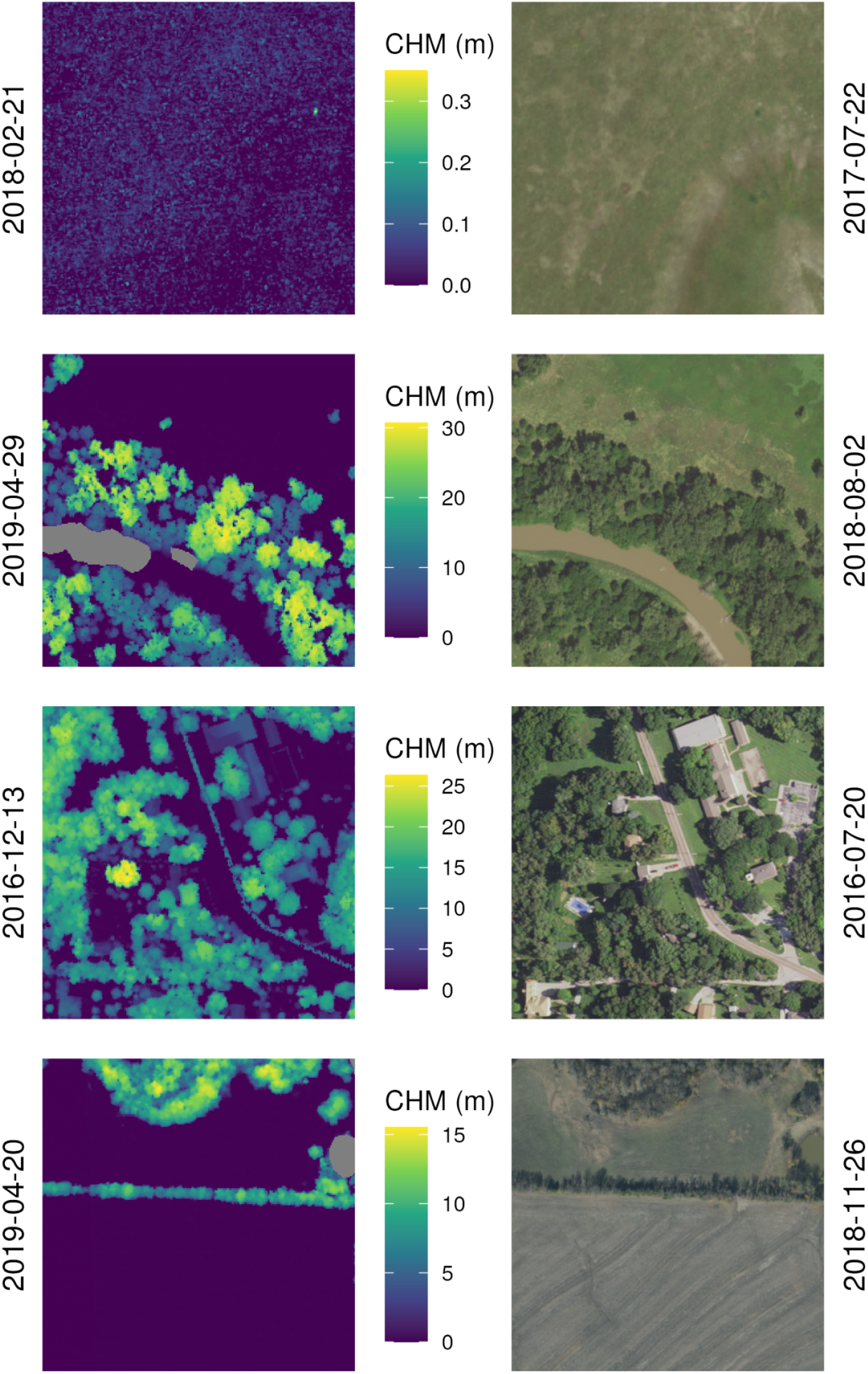
A sample of paired canopy height model (CHM; left) and NAIP images (right). Dates on vertical axes represent the date of acquisition.

**Figure 5.**
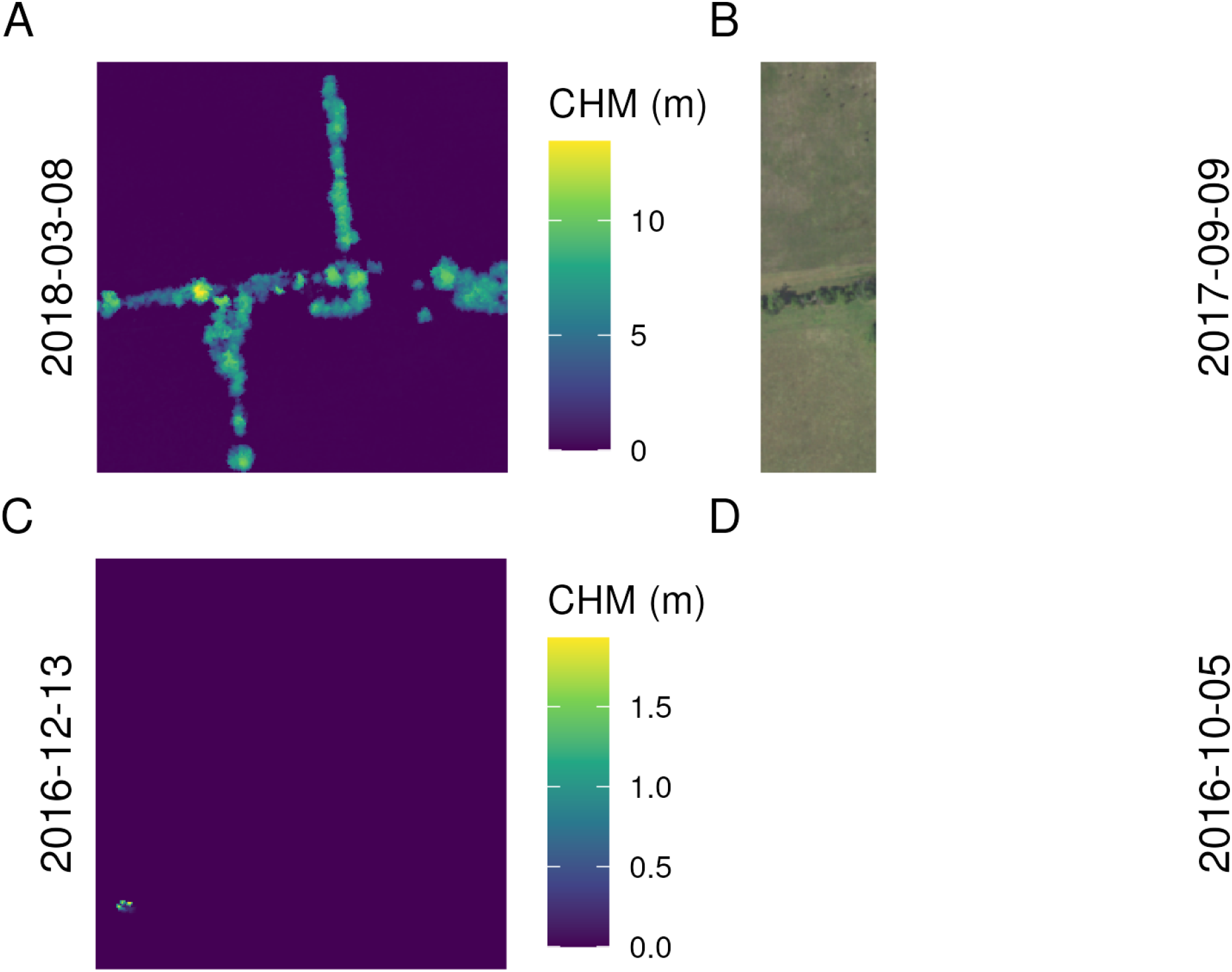
Canopy height model (CHM; left) and NAIP images (right) where the NAIP image is partially (B) or completely (D) masked. A “mask” band is supplied with each NAIP image. Dates on vertical axes represent the date of acquisition.

**Figure 6.**
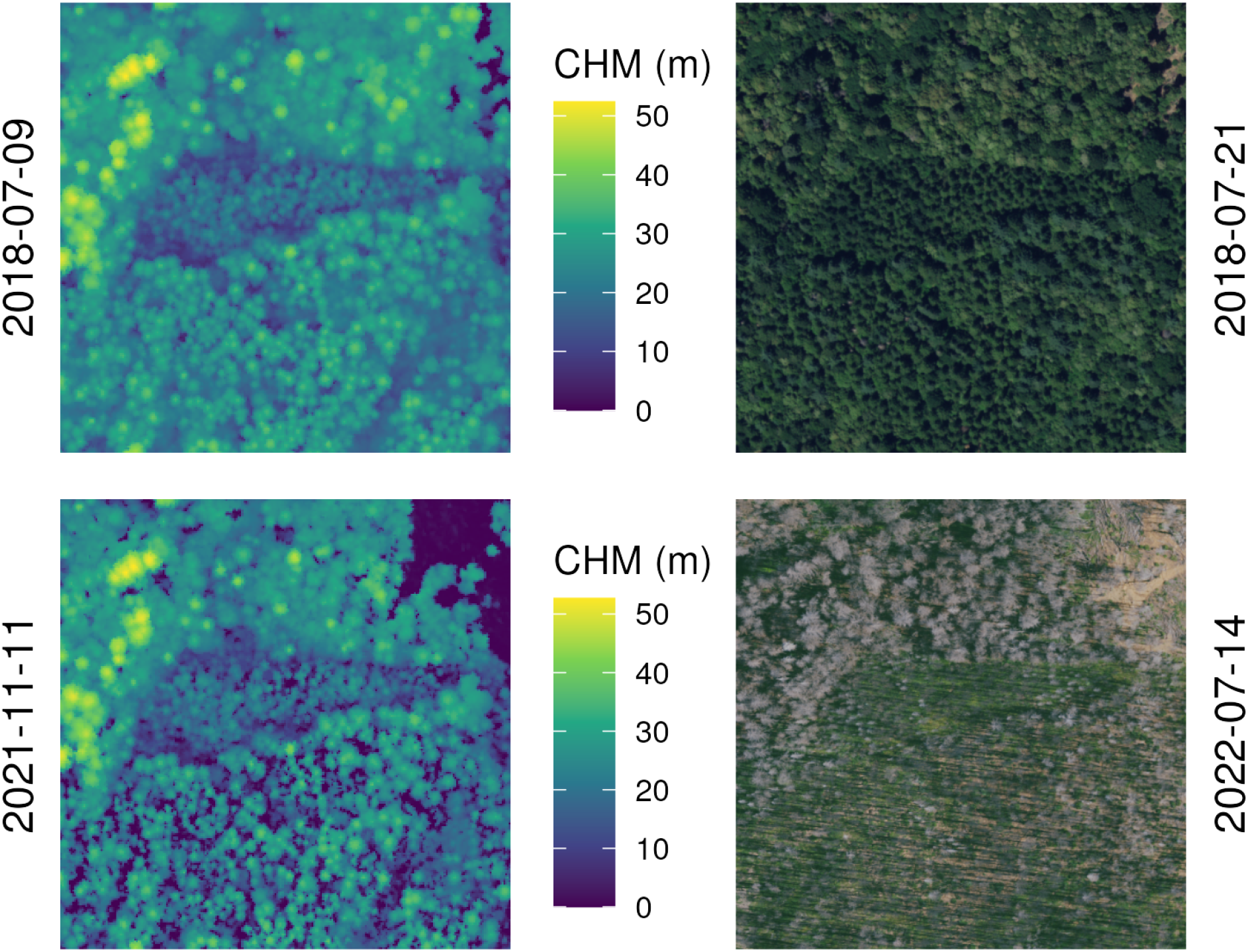
Canopy height model (CHM; left) and NAIP image (right) of the same sampling location with multiple CHM and NAIP acquisitions. Note the differing features between the CHM and NAIP image in the bottom acquisitions. Dates on vertical axes represent the date of acquisition.

We compared the output of our methodology against CHMs produced by the National Ecological Observatory Network (NEON). NEON releases their lidar and CHM data in 1km × 1km tiles^18,19^. NEON produces their CHMs with a different methodology, using LASTools and an implementation of the Khosravipour et al. pitfree algorithm^16,19^. We retrieved a random tile from four random NEON sites and produced a CHM using our methodology. We evaluated our CHM by calculating root mean square error (RMSE), mean absolute error (MAE), and the coefficient of determination (r^2^) against the NEON produced CHM. Error metrics and data distribution (Table 2; Fig. 7), as well as visual inspection (Fig. 8), revealed comparable outputs.

**Table 2.**
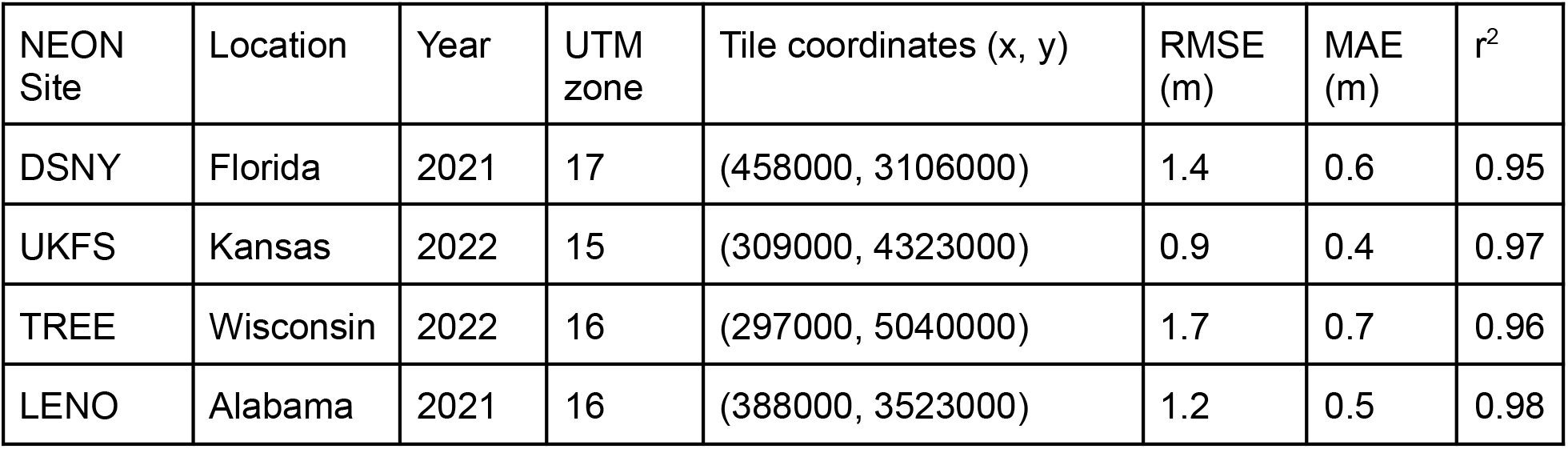
Comparison metrics (root mean square error, RMSE; mean absolute error, MAE; coefficient of determination, r^2^) of canopy height model estimates for four random NEON sites. Canopy height model estimates were produced using the methodology in this paper and compared to NEON produced canopy height models.

| NEON Site | Location | Year | UTM zone | Tile coordinates (x, y) | RMSE (m) | MAE (m) | $r^2$ |
| --- | --- | --- | --- | --- | --- | --- | --- |
| DSNY | Florida | 2021 | 17 | (458000, 3106000) | 1.4 | 0.6 | 0.95 |
| UKFS | Kansas | 2022 | 15 | (309000, 4323000) | 0.9 | 0.4 | 0.97 |
| TREE | Wisconsin | 2022 | 16 | (297000, 5040000) | 1.7 | 0.7 | 0.96 |
| LENO | Alabama | 2021 | 16 | (388000, 3523000) | 1.2 | 0.5 | 0.98 |

**Figure 7.**
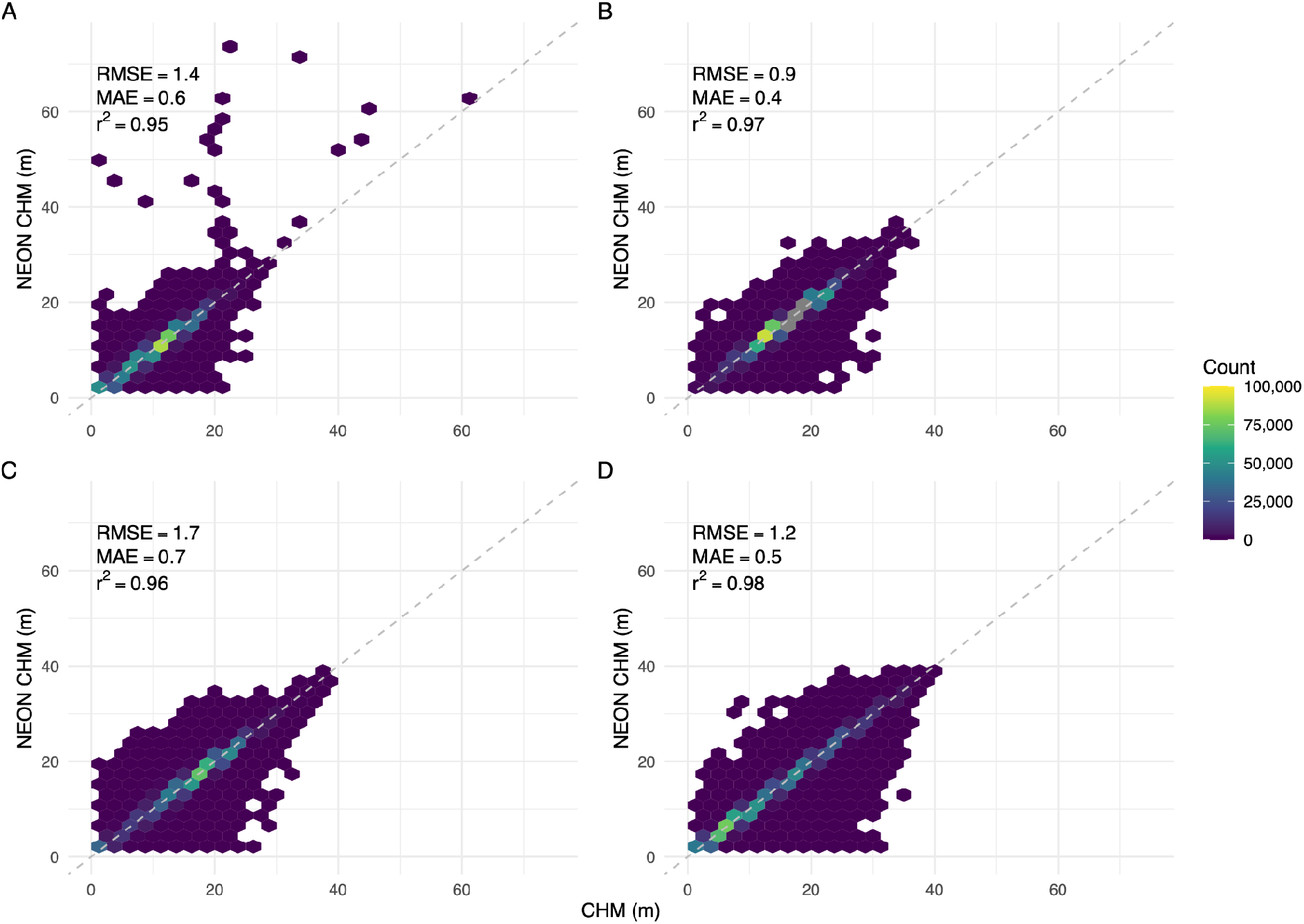
Canopy height model (CHM) estimates (m) produced using the methodology described herein (horizontal axis) relative to estimates produced by the National Ecological Observatory Network (NEON; vertical axis) for four NEON sites: A) DSNY; B) UKFS; C) TREE; and D) LENO. The dashed gray line represents a 1:1 relationship. See also Table 2.

**Figure 8.**
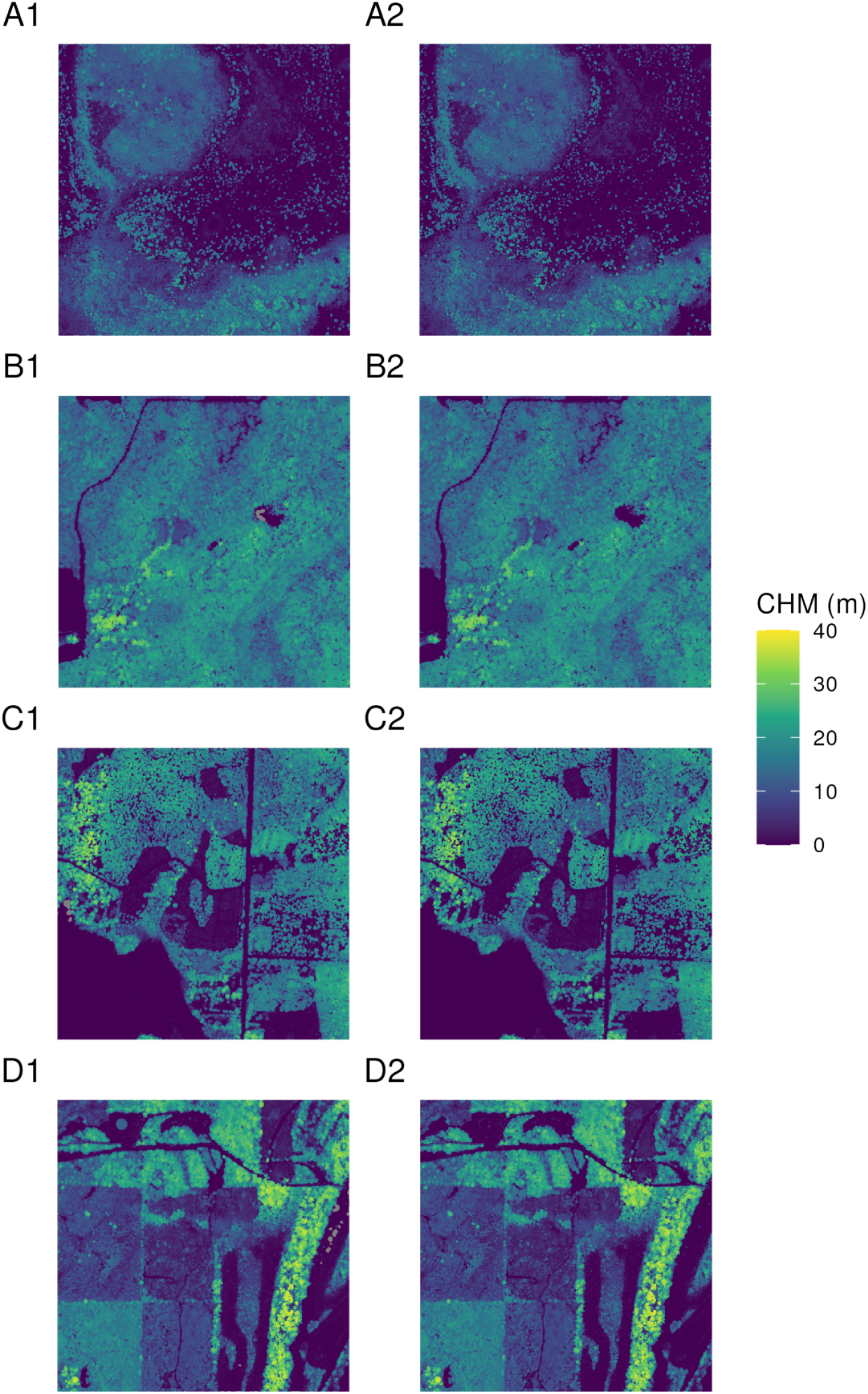
Canopy height model (CHM) estimates produced using the methodology described herein (left column) and estimates produced by the National Ecological Observatory Network (NEON; right column) for four NEON sites: A) DSNY; B) UKFS; C) TREE; and D) LENO. The spatial extent of each image is 1km × 1km.

## Code Availability

Code used to retrieve and process lidar data and NAIP imagery are available on GitHub (https://github.com/allredbw/chm-naip).

## Acknowledgements

This research used computer resources provided by 1) the SCINet project and/or the AI Center of Excellence of the USDA Agricultural Research Service, ARS project numbers 0201-88888-003-000D and 0201-88888-002-000D; 2) Google Earth Engine; and 3) the Numerical Terradynamic Simulation Group at the University of Montana. Any use of trade, firm, or product names is for descriptive purposes only and does not imply endorsement by the U.S. Government.

## Author information

### Authors and affiliations

**Numerical Terradynamic Simulation Group, University of Montana, Missoula, MT, USA** Brady Allred and Scott Morford

**Jornada Experimental Range, USDA Agricultural Research Service, Las Cruces, NM, USA** Sarah McCord

### Contributions

Allred performed data retrieval and processing, and wrote the manuscript. McCord and Morford provided fantastic intellectual guidance and assisted with the manuscript.

